# Infant vocal category exploration as a foundation for speech development

**DOI:** 10.1101/2024.01.17.576142

**Authors:** Hyunjoo Yoo, Pumpki Lei Su, Gordon Ramsay, Helen L. Long, Edina R. Bene, D. Kimbrough Oller

**Affiliations:** Department of Communicative Disorders, College of Arts & Sciences, The University of Alabama, Tuscaloosa, Alabama, United States of America; Department of Speech, Language, and Hearing, School of Behavioral and Brain Sciences, The University of Texas at Dallas, Richardson, Texas, United States of America; Spoken Communication Laboratory, Marcus Autism Center, Children’s Healthcare of Atlanta, Atlanta, Georgia, United States of America; Department of Pediatrics, Emory School of Medicine, Atlanta, Georgia, United States of America; Waisman Center, University of Wisconsin–Madison, Madison, Wisconsin, United States of America; Origins of Language Laboratory, School of Communication Sciences and Disorders, University of Memphis, Memphis, Tennessee, United States of America; Institute for Intelligent Systems, University of Memphis, Memphis, Tennessee, United States of America; Konrad Lorenz Institute for Evolution and Cognition Research, Klosterneuburg, Austria

## Abstract

Non-random exploration of infant speech-like vocalizations (e.g., squeals, growls, and vowel-like sounds or “vocants”) is pivotal in speech development. This type of vocal exploration, often noticed when infants produce particular vocal types in clusters, serves two crucial purposes: it establishes a foundation for speech because speech requires formation of new vocal categories, and it serves as a basis for vocal signaling of wellness and interaction with caregivers. Despite the significance of clustering, existing research has largely relied on subjective descriptions and anecdotal observations regarding early vocal category formation. In this study, we aim to address this gap by presenting the first large-scale empirical evidence of vocal category exploration and clustering throughout the first year of life. We observed infant vocalizations longitudinally using all-day home recordings from 130 typically developing infants across the entire first year of life. To identify clustering patterns, we conducted Fisher’s exact tests to compare the occurrence of squeals versus vocants, as well as growls versus vocants. We found that across the first year, infants demonstrated clear clustering patterns of squeals and growls, indicating that these categories were not randomly produced, but rather, it seemed, infants actively engaged in practice of these specific categories. The findings lend support to the concept of infants as manifesting active vocal exploration and category formation, a key foundation for vocal language.

## Introduction

### Clustering of vocal types in early human development

To uncover mechanisms underlying language acquisition and the distant origins of language, various learning mechanisms have been proposed, including imitation and social interaction [1, 2]. Infants’ endogenous vocal production and exploration of vocal types has received comparatively little attention in the literature until recently [3]. The present study provides empirical evidence of the early emergence of vocal development in infants by systematically quantifying the occurrence of infant vocal categories, suggesting that active vocal exploration is fundamental to subsequent speech development and provides a foundation for interaction with caregivers.

The work is inspired by the fact that various vocal types of early human vocalizations known to be precursors to speech (“protophones”) have been observed to occur in clusters [4, 5] of particular phonatory types. There are three predominant types: 1) vocants are vowel-like sounds produced with normal phonation in the mid-pitch range of the individual infant; 2) squeals are produced at very high pitch, often in falsetto phonation; and 3) growls are low pitched or harshly phonated sounds often in fry. Clustering can be observed in that a baby may produce several squeals in a short period of time, and then for the next few minutes no squeals at all, while other vocal types continue to be produced. Also, from day to day there appear to occur wide variations in the number of squeals babies produce. Similar clustering has been observed for growls and other less frequently occurring sounds such as raspberries or whispers [6]. In the present paper we will greatly expand the quantitative exploration of this presumable non-randomness of infant vocal types by reporting on a sample of 130 infants recorded longitudinally all-day in their homes. In the present paper we define clustering as the non-random occurrence of particular protophones across sessions, that is, across specified time periods. The paper does not address immediate “repetition” of particular protophone types, one after another within sessions.

The pattern leads us to consider why a tendency for clustering exists. The activity often seems playful, and one cannot avoid wondering if it might constitute practice. But practice for what? If for language, then we are forced to ask how being capable of producing such sounds as squeals and growls could form foundations for language, because such utterances are not in and of themselves elements of language. They are not well-formed syllables, and they do not, indeed they cannot become words. We are driven to question why there could be any advantage to practicing such sounds. Our paper is based on the supposition that clustering of infant vocal types 1) helps to establish the principle of vocal category formation, the mastery of which is required for vocal language [6] and 2) provides information to caregivers about infant wellness.

### Foundations for mature vocal communication in infancy

In the first year of life, human infants engage in remarkably extensive vocal activity with protophones, a term that includes both precanonical and canonical speech-like sounds, while excluding cries, laughs and vegetative sounds. This vocal activity has been estimated to involve an average of 4-5 protophones per minute every waking hour from the first month of life and continuing throughout the first year [3]. Caregivers respond to protophones in face-to-face sustained interactions called “protoconversations” [7, 8] that have no precedent in the animal kingdom as far as we know.

Surprisingly, even though these salient interactive periods between human caregivers and infants have generated considerable interest and speculation about foundations for language, the great bulk of the protophone production of human infants does not actually occur during social interaction. Instead, examination of extensive laboratory recordings along with all-day recordings in infant homes has led to the conclusion that more than 90% of protophones are produced in endogenous, non-socially-directed activity [9], much of which might be termed infant “vocal exploration” or “vocal play” [5]. In an attempt to understand the mechanisms of language development, researchers have explored cross-species “babbling” in non-human primates and songbirds (for more information, see [10–14]). However, active vocal exploration seems to be uniquely observed in human infants, distinguishing them from other species. In all the cases where birds, bats, or humans engage in seemingly playful infant vocal activity, it is presumed that the babbling is a precursor to the mature form of vocal communication, and that the activity can be thought of as a kind of exploratory practice laying groundwork for the elaborate forms of song and speech.

### Categories of early vocalization and the idea of infant vocal practice

Widely accepted descriptions of early human vocal development designate categories of phonation that appear to occur universally in the first months of life [15–18]. Although they can be broken down into further subcategories, the three most common phonatory categories include:

1) vocants (also called vowel-like sounds) produced in normal phonation, the kind that dominates the utterances of natural languages, and produced at the mid-pitch range of the individual, 2) squeals, which are saliently high-pitched sounds, produced with typically at least twice the fundamental frequency (f_0_) of the infant’s usual voice, and 3) growls, produced typically with salient harsh, noisy phonation, often at low pitch, or produced in vocal fry (“pulse”) at very low pitch with respect to the infant’s typical f_0_ [19]; generally, no sound that is deemed to occur at higher than the mid-pitch range for the individual is categorized as a growl.

These three vocal types have been reported to occur not only by ethologically oriented observers who have tracked infants with longitudinal recordings [5, 16, 20], but also by parents who have often responded to open-ended questions about their infants’ vocal sounds in the first half year of life with terms drawn from the common parlance to describe them, in English, vowel-like sounds (which in our technical terminology are called vocants), squeals and growls. Other languages also often have terms to designate the primary phonatory protophones [6].

Of course, additional infant sounds do occur (raspberries, ingressive sounds, whispers, and so on) though far less frequently, and many utterances combine phonatory characteristics of the three primary protophones. As the first year progresses, infant utterances begin to show more speech-like characteristics, and by the second half year, more complex syllables occur, culminating in canonical babbling, with well-formed syllables such as “ba”, “da”, or “na”, and often reduplicated sequences of these sounds (“baba”, “nanana”, and so on) [21–23]. The phonatory characteristics of vocants, squeals and growls persist throughout the first year, and even the most speech-like fully canonical sequences are produced with phonatory properties that allow them to be categorized as vocants, squeals and growls, just as in the case of precanonical protophones.

Research having tracked the occurrence of protophones has long posited that vocal play, in particular with the three primary phonatory types, involves clustered production of the individual types, a non-random occurrence that has suggested infants may be engaged in vocal practice, perhaps in an attempt to consolidate the categories themselves [16, 22]. This reasoning is founded in the idea that the categories emerge from exploration, rather than being innate vocal givens or learned from parents. The three principal protophones are thought to be creations of the infant, and the fact that they seem to fall into particular categories is thought of as the result of exploration in a non-random landscape of phonatory possibilities, where Waddingtonian wells of attraction [24] tend to draw the exploratory activity into the three phonatory types. The pattern of development is reminiscent of the idea of quantal categorization in speech perception [25].

The fact that parents seem to notice the distinctions among the three has been proposed to be attributable at least in part to their non-randomness of occurrence across time [6]. An infant may be heard producing a train of squeals, for example, and although the great majority of protophones are vocants, the parent may notice the squeals as salient departures from the more typical vocants especially if they tend to be repeated or to occur in concentrated clusters. Parents engaging in face-to-face vocal interaction with their infants often attempt to elicit one of the three phonatory protophones by producing imitated versions of the sounds they believe to be in the infant repertoire. It is as if parents seek to confirm their infants’ emerging vocal competence in a game of mutual imitation, where parents most often initiate imitation by trying to draw infants into producing one of the protophone types already under their command [26].

So the non-random occurrence of the protophone types is seen as an important feature of infant vocal development in two ways. First, clustering may constitute a type of practice, where infants seek to firm up their control of vocal categories through intentional manipulation of each type. This manipulation and learning of new types appears to constitute a critical foundation for language, because one of the necessary requirements of language is the ability to adapt to using new vocal categories (intonational types as well as syllables and phonemes) that are language specific and to provide the basis for learning to use indefinitely large inventories of possible words and sentences [6]. Second, if the reasoning is correct, clustering provides a basis for caregivers to recognize infant progress toward language in the sense that they can thereby discern infant category control. This parental discernment would seem to begin with simple recognition of the primary phonatory protophones, a recognition forming a basis for elaborate vocal interchanges with infants. Later in the first year, parents recognize the appearance of repetitive canonical syllables, which results in an intuitive reaction whereby parents begin to “negotiate” with infants [27] over the possible meanings of particular syllable sequences (yes, you said ba, it’s a ball, say ba…) [28].

### Why has there been so little research on vocal practice by clustering in human infancy?

Given the seeming importance of clustered production of vocal types in infants, it may be seen as odd that very little research has been devoted to quantitative determination of the extent of such practice or its course of development. As far as we know, only we and some of the collaborators of the University of Memphis Origin of Language Laboratories (OLL) have addressed the issue empirically and in each case at small scale. 1) In the Supplementary Material to one of our papers [4], we presented a demonstration using lag sequential analysis [29, 30] of a tendency of 9 infants in brief laboratory recordings to produce the three primary phonatory types in non-random sequences where the identity of an infant utterance (for example, a growl) predicted the likelihood that the next utterance in sequence would be of the same type at a higher than chance level. 2) In a separate demonstration reported in the same Supplementary Material, recurrence quantification analysis [31] was used to illustrate the clustering of squeals in a sequence of utterances from an infant, that is, a tendency for few squeals to be present at the beginning of a particular recording session, but a large number of squeals to cluster at the end of the session. 3) In a separate paper [32], we reported “session effects”, whereby 3 infants showed a tendency to produce durational patterns in babbling that were notably different from one period (or session) of recording to another for the same infant, even within the same day.

Such session effects have been recognized throughout decades of longitudinal research on vocal development in infancy by the OLL, having required us to conclude that small-scale sampling by recording of infant vocalizations could not be expected to yield representative samples of any aspect of infant vocal patterns (see also [33]). From one period to another, a wakeful and alert infant can change from being completely silent to being very voluble, and during periods of volubility [34], from producing a very limited repertoire of vocal types to producing a wide array of types. With a comfortable baby, the most common protophones by far are vocants [4]. But we have long noted that periods occur where squeals or growls can abruptly take center stage, and when they do, an observant individual cannot help but take notice.

There are several reasons that vocal exploration, particularly the quantification of vocal clustering, has received limited attention in the study of infant vocal development. First, researchers have historically focused heavily on the anatomic and physiological aspects of infant vocal development, viewing the vocal products as showing gradual progression from immature and unstructured forms to mature speech [35–39]. While this perspective has merit, it tends to overlook the critical role of infants as exploratory agents in their own vocal development. This perspective has been influenced by longstanding views, from the concept of the “tabula rasa” in Western philosophy to Jakobson’s claim that prelinguistic vocalizations are random byproducts of biological inclinations [40]. These views are outdated, yet the tendency to underplay infant active involvement in vocal development persists.

Second, while some scholars have recognized and described vocal exploration in infancy, the precanonical stage of phonatory control development has received considerably less attention than the canonical stage and higher-level language development (e.g., words and sentences) in childhood (see review in [41]). Yet without the fundamental capability of phonatory control, articulatory development as manifest in canonical babbling would seem largely superfluous, since syllables overwhelmingly require phonation in their nuclei, and articulatory movements without phonation are largely soundless.

Third, in terms of the mechanisms of language development, social interaction and imitation have been the primary focus in the literature, with the seeming assumption that infants learn vocal categories by listening to parents, interacting with them vocally and imitating the existing baby sounds [42–44]. But in fact, only a small proportion of infant vocalizations are produced during vocal interaction, and imitation of parental sounds is extremely rare in early life [26, 45]. In fact, it appears that the vast majority of apparent imitation by infants may actually be based on parental elicitation of the sounds parents know the infants already have in their repertoires, for example, squeals, growls, and vocants [28]. It is difficult to find evidence that infants introduce any new sounds into their exploratorily developed vocal repertoires as a result of listening to parental talk until late in the first year, when the first words begin to appear.

Lastly, most prior studies on infant vocal development have been conducted in a laboratory, for only a few minutes to about an hour [2, 46, 47]. Therefore, vocal exploration may often not have been salient enough to capture researchers’ attention during such constrained time periods, and especially since parents are often instructed to elicit vocalization from their infants during such recordings. However, with the widespread availability of all-day home recordings in recent time [48–50] more representative data are now being obtained from naturalistic environments.

### Purposes of the present study

Our goal is to provide a large-sample quantitative assessment of the tendency of typically developing human infants to produce the three primary protophone categories in clusters, rather than simply randomly distributing the categories across segments of time. This assessment, taking advantage of a massive database of recordings described in Methods, will also offer perspective on the degree to which infants across the first year give evidence of vocal category development, which is a crucial capacity in language. The work should also provide perspective on vocal practice in infancy.

The present study targets two primary questions: 1) to what extent do infants tend to produce phonatory categories in clusters rather than randomly across segments in recordings, and 2) to what extent do clustering patterns change with age during the first year of life.

## Method

### Participants

A total of 130 typically developing, English-learning infants participated in the present study (55% were male and 63% were designated as living in homes of relatively high SES as indicated by maternal education level). Participants whose data were analyzed in this study were recruited from September 2012 to October 2020. These infants were recruited in the Atlanta area as part of a much larger longitudinal study conducted by the Marcus Autism Center, Children’s Healthcare of Atlanta and Emory University School of Medicine (NIH P50 MH100029) investigating vocal development in infants who 1) had no family history of autism or other developmental disorders or 2) had an older sibling with confirmed diagnosis of autism and thus were at elevated likelihood for autism. All 130 infants included in the present study were classified as having no clinical features at 2 and 3 years after thorough evaluations by Marcus Autism Center expert clinicians. Among the 130 infants, 103 were classified at enrollment as being at low likelihood for autism diagnosis. We report data on all 130 after having verified that the 27 at elevated likelihood showed very similar proportions of recordings with significant growl and squeal clustering to the low likelihood group of 103.

The OLL contributed human coding of the recordings of the infants as part of a collaborative NIDCD longitudinal study between the University of Memphis and the Marcus Autism Center/Emory University (NIH R01 DC015108). All the recording/coding protocols and consent documents signed by all the parents were approved by the Institutional Review Boards (IRBs) of Emory University and the University of Memphis. The third author at Emory University had access to identifying information for all participants in the present study. The data for the present work are not secondary. The recordings were collected under the supervision of the third author in Atlanta, and the coding was conducted under the supervision of the last author in Memphis.

### Recordings and the selection of 5-minute segments for coding

Language ENvironment Analysis (LENA) all-day recorders [51] were used to collect the audio data. The recorders are small and light enough (about the size of an iPod) for infants to carry in a pocket of a vest without any disturbance. Because LENA allows recording up to 16 hours a day with high audio quality (16 kHz sampling rate), it enables researchers to obtain representative data on infant vocalization and the auditory environment in the home. The recorder has been used routinely in a wide variety of research on early language development, starting with Zimmerman et al. [48].

Caregivers were instructed as in a variety of our prior studies on how to use a LENA device, including how to start, stop and pause recording. All recordings were carried out remotely in the home, once every month from birth to two years of age, and were scheduled as far as possible on the same calendar day of each month to ensure rotation of weekdays.

Recordings were mailed out and retrieved via USPS priority mail. Each completed recording was uploaded through the LENA software to save the data. For further recording details see [52] for a description of the procedures that were used at the Marcus Autism Center, where the recordings were conducted.

A total of 1154 all-day recordings were collected, an average of 8.9 recordings per infant (range 4-12). The recordings were always available for coding in the OLL and have been accessed repeatedly from September 1^st^, 2015 to the present. The human coding was conducted on 21 randomly selected 5-minute segments from each of the recordings. At the point of analysis, each available recording was assigned to one of six age groups as follows: 0-2 months, 3-4 months, 5-6 months, 7-8 months, 9-10 months, and 11-13 months.

### Coders, training, and coding environment

Thirty-six English-speaking graduate students in the University of Memphis School of Communication Sciences and Disorders were trained and coded the recordings. They were trained in phonetic transcription in their program of study but were more intensively trained for infant vocalization research by the fifth and the last author regarding all the coding parameters, including the primary phonatory types (i.e., squeals, growls and vocants). The six- to eight-week training for the coding of infant vocalizations is described in greater detail in some of our prior work [3, 52]. The coders met periodically across the training period for weekly one-to-two-hour lectures and presentation of audio examples drawn from the OLL archives of infant vocalizations. After each lecture session, they conducted practice coding on recording segments. At the end of the week, these practice sessions were reviewed with the trainers during the lecture sessions and as needed individual coders had individual sessions with the trainers to address discrepancies between the categorizations that had been made by the trainees and also by key codes that are available for many of the segments based on coding by the last author, who is the longest term researcher on vocal development among any of the OLL participants. Coders were required to meet standards of agreement with the key codes (to be within 10% of the counts on protophones) by the 6^th^ week of training or else additional training would ensue.

As a part of the training and providing a basis for coder agreement research on the three phonatory protophone categories, all the coders completed some practice coding on several of 9 full recordings where they coded all 21 randomly selected 5-minute segments independently of any other coder. These were not the research recordings on the 130 infants but were drawn instead from the same archives of LENA recordings at the Marcus Autism Center that produced the data on the 130 infants. These recordings are part of a set of recordings from the Marcus Autism Center utilized in the OLL for training of coders and about which agreement data are presented below.

Once trainees had completed training, they were admitted to the “coding team”, which typically consisted of about 12 individuals year-by-year, with a new group of trainees being recruited and trained in the fall semester of each year, replacing individuals having graduated that spring or summer.

A total of 24,234 5-minute segments were coded. The coders were assigned to groups of four infants, being blind to the age, risk status, and diagnostic outcome of the infants. They coded each recording completely (all 21 randomly selected segments in chronological order as occurring in the recordings from morning to night time) before proceeding to the next recording, which was randomly selected from among all the recordings from four infants assigned to each coder. Many of the coders were assigned to additional groups of four infants depending on their availability during the period of the research. The order of infants whose recordings were to be coded by each coder was also randomly assigned. Each individual coded all the recordings of an average of 4 infants (range 1 to 9 infants per coder).

Action Analysis, Coding, and Training (AACT [53]) was used for identification of each vocal type. AACT is a software environment enabling coders to label any kind of action in audio or video or both (see [4] Supplementary Material for details). Coders used keystroke or mouse-selected coding for each of the three protophone categories (squeals, vocants, growls), along with a few additional very infrequently occurring protophone categories (ingresses, whispers, raspberries, and non-phonated frication sounds) plus cries, whimpers, and laughs, while listening to each of the 5-minute segments. Coding was conducted in real time, so that each 5-minute segment took 5 minutes to code. Each coder responded to a brief questionnaire at the end of each coding segment, adding about another minute to coding time per segment. They answered the questions on a Likert-like scale from one to five in order to provide additional relevant information for each segment. For the present study, the key question concerned infant sleep. If the coder determined the infant to be asleep during the whole 5-minute segment, a value of 5 was assigned to that question, and the segment was then not included in the clustering evaluation for that recording.

### Coding categories

The coding yielded counts of squeals, vocants, and growls, leaving aside many other possible vocalizations of infants that occur far less frequently and are not considered to be precursors to speech. All coding was conducted at the utterance (that is, the breath-group) level. Extensive additional coding category information as well as details on how squeals, vocants, and growls were defined, along with spectrographic illustrations, are provided in Supporting Information, in the initial section called Coding categories, definitions and coding criteria.

### Data processing and statistical analysis of clustering

#### Hypotheses tested

The 21 randomly selected 5-minute segments coded for each available recording for each infant were tested for clustering of squeals with respect to vocants in one case and growls with respect to vocants in the other. In addition, the data were binned into six age groups to evaluate the possibility that clustering patterns might change with age.

#### Preliminaries to data analysis

We applied several exclusion criteria for segments prior to conducting Fisher’s exact tests to determine evidence of clustering. For example, segments with high cry or whimper rates or segments where the infant was asleep were assumed to be incompatible with vocal play. Thus, we excluded segments where 1) 5 or more negative (cry and whimper) utterances occurred or 2) the infant was asleep throughout the 5 minutes according to the coder questionnaires. We also designated as not analyzable (NA) for the squeal vs. vocant comparison any recording where the sum for squeals over all segments was 0 (i.e., no squeals were coded), and similarly, we designated as NA for the growl comparison any recording where the sum for growls was 0 (i.e., no growls were coded). Segments remaining after these exclusions were termed “surviving” segments.

Finally, we designated as NA any recording that had only 1 surviving segment because at least 2 segments are required to conduct Fisher’s exact tests. Those requirements having been met, a significant Fisher’s exact test for the available comparisons within a recording would indicate that the protophone types were distributed in a non-random way, indicating clustering. In other words, either squeals or growls or both were being produced significantly more often in particular segments than in other segments of the same recording. Tables S1 and S2 (in Supporting Information, Illustrations of the application of Fisher’s exact test) present examples from the real data of application of the Fisher’s exact test for recordings, where the comparison is between squeal and vocant counts in S1 and between growl and vocant counts in S2. These illustrations are designed to help the reader understand the analysis method intuitively.

#### Statistical analysis

Fisher’s exact test was selected as an appropriate test of whether the proportions for one variable (types of vocal categories) were different with respect to values of the other variable (5- minute segments) across each recording. This test is particularly suitable for our data given that the raw count of each vocal category was often small and was usually uneven across cells (i.e., segments). In our case, we sought to determine whether the proportions for squeals and growls were different across 5-minute segments in a recording with respect to the proportion of vocants. The null hypothesis was that the proportion of squeals/growls or vocants was the same with respect to vocants across segments. A significant result indicated there was a significant difference in squeals/growls and vocants across segments, and thus that squeals/growls did not distribute randomly with respect to vocants.

### Coder agreement

To establish a foundation for assessing the reliability of coding across three protophone types, we conducted a comprehensive evaluation of coder agreement through various analyses (for details, refer to Supporting Information, Coder agreement). Our evaluation focused on a very large coding agreement set, comprising 21 segments from each of the 9 recordings, coded during the training period. The agreement set enabled us to obtain a very large number of intercoder agreement pairings because, on average, about 32 coders independently coded the same 21 segments for each recording. A second, though smaller, agreement study involved 9 coders, which was conducted toward the end of the coding by those individuals on 523 segments semi-randomly selected from the data reported in Results. It makes sense to assume coder agreement would be lower during the training period and higher after actual data collection. The higher agreement obtained during the second, though smaller evaluation, tends to confirm this assumption.

These evaluations including coefficients of variation and Spearman rank order correlations demonstrated that coder agreement was highly statistically significant (see Supporting Information, Coder agreement). Permutation tests conducted on the 9 recording agreement set further confirmed that coder agreement was more than adequate to justify the analyses of clustering reported below.

## Results

We cleaned the data to ensure they met our exclusion criteria in accord with the specifications above, before conducting Fisher’s exact tests. Thus, at the outset of the analysis, we identified the segments where infants were asleep according to the coders. Among all the recordings, 18,399 segments (76%) were identified in which the infants did not sleep, according to the coders’ questionnaire responses, throughout the entire 5-minute period. We then located the number of segments that included too many negative vocalizations, defined as the sum of cries and whimpers being greater than or equal to 5. Fourteen percent of the non-sleep segments were thus eliminated from consideration in the Fisher’s tests. These eliminations left 15,774 surviving segments for potential evaluation. Finally, we evaluated all the recordings for the possibility that they had 0 squeals or 0 growls. There were 125 recordings with no coded squeals and 123 with no coded growls. These recordings were designated as NA and were not evaluated by Fisher’s exact tests on squeals vs. vocants or growls vs. vocants respectively.

### Percentage of recordings determined to show clustering

There was a considerable tendency for the infants to show significant clustering patterns.

For squeals, 40% of the recordings showed significant clustering (*p* < .05) by the Fisher’s test, and for growls the value was 39% (*p* < .05). These percentages of recordings that met the significant clustering criterion for each infant at each age were computed to include the NA recordings (which of course were not analyzed by Fisher’s test) in their denominators, and consequently they supply an indication of squeal and growl clustering across *all* the recordings available for each infant. Since we only selected 21 segments from each all-day recording for coding (only about 15% of the available recording time), these values surely underestimate the percentage of days the infants actually engaged in some clustering of squeals and/or growls. The data thus estimate a lower bound on the amount of clustering of squeals and growls at around 40% of all-day recordings.

Furthermore, evaluating whether individual infants showed *either* significant squeal clustering *or* significant growl clustering on each recording, we found that 61% of recordings showed a significant amount of clustering for one or the other. So it can be concluded that even though we sampled randomly from a relatively small proportion of the day, infants *usually* showed some discernible clustering activity of protophones.

At the infant level, we evaluated all the recordings available for each infant and found that 87% of infants showed at least one age at which their recordings showed significant squeal clustering *and* at least one age with significant growl clustering. There was not a single infant who, on evaluation of all the available recordings for the infant, showed neither a significant case of squeal clustering nor of growl clustering. Only 8 infants (6%) had 3 or fewer recordings with significant squeal or growl clustering. In contrast, 35 infants showed 6 or more recordings with significant growl clustering and 30 showed 6 or more with significant squeal clustering. Some infants seemed clearly to cluster growls more than squeals (about 10% of infants showed at least 3 more recordings with significant growl clustering than squeal clustering) while some (about 8%) clustered squeals more than growls.

### Percentage of recordings with significant clustering across ages

When the data were analyzed by age for each vocal type comparison, we found significant clustering patterns of vocal types across all age groups. Fig 1a-1c supply a summary of the data, indicating that clustering occurred at all the ages. The error bars represent 95% bootstrapped confidence intervals computed around the means for all infants at each age level. Infant data were averaged for all the recordings of each infant within the age interval prior to computing means and confidence intervals.

**Fig 1.**
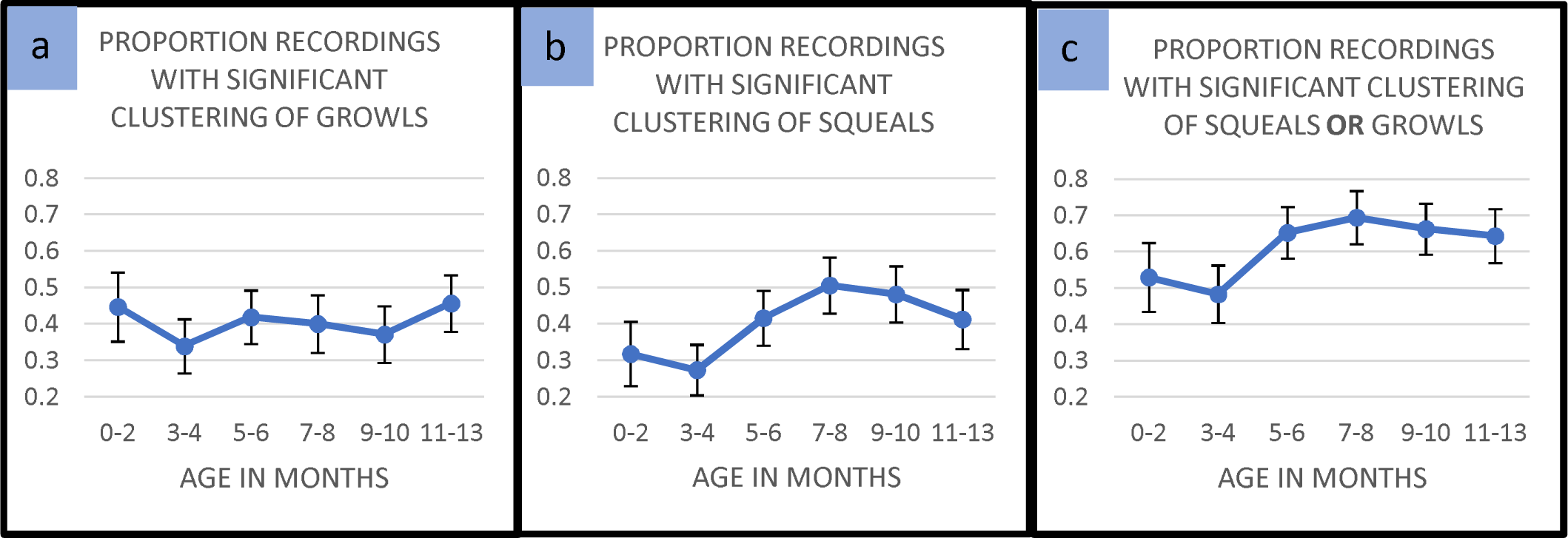
Clustering across Ages: The figure shows means of statistically significant clustering for recordings from infants at each of 6 age intervals. Values were computed at the infant level, such that all recordings for any infant within the age intervals indicated were averaged first, and 95% bootstrapped confidence intervals displayed as error bars were computed based on the averages for the number of infants who had recordings within those age intervals. **a.** Growls showed significant clustering for more than 30% of the infants at all ages. **b.** Squeals showed significant clustering for more than 30% of infants at all ages except the 3–4-month interval, where 27% of infants had significant clustering of squeals. **c.** In the final panel we display the proportion of infants who showed either significant growl or squeal clustering at each of the age intervals.

In panels 1a and 1b the data show considerable occurrence of squeal and growl clustering in the very first months of life. Interestingly, the highest amount of clustering did not fall within the 3–4 month range, traditionally thought of as the period for vocal play in stage models of vocal development [16]. In fact, the 3–4-month interval showed the *lowest* mean values for both squeal and growl clustering. Squeal clustering showed a tendency to increase toward the middle of the year, peaking at 7–8 months. The squeal clustering pattern appeared to vary more across age than the growl clustering pattern, but in both cases, the final interval of the year, like the first interval, showed substantial clustering. Thus, a pattern of clustering occurred at all ages.

From 48% to 69% of infants showed significant clustering of either growls or squeals at the various age intervals, and all the age intervals beyond 5 months revealed more than 60% of infants had significant clustering.

In Fig 1c, the data are presented in such a way that either squeal clustering or growl clustering was counted. Thus, each data point represents the proportion of infants whose recordings at the age in question showed either significant growl or squeal clustering. Consequently, the means are higher at each age than for either Fig 1a or 1b, illustrating the fact that infants often tended to produce recordings with clustering of squeals or growls but not both.

The confidence intervals supply information about possible significant differences across the ages. The apparent differences suggest, for example, a possible tendency for clustering overall (panel 1c) or clustering of squeals (panel 1b) to be more common beyond 5 months of age than before 5 months. Yet there are important provisos to offer about these apparent age differences, as we explain below.

## Discussion

It has long been observed, mostly without quantification, that human infants produce vocalizations in systematic patterns where some protophone categories occur in clusters across time, and it has been thought that these patterns of non-random occurrence might represent practice of categories emerging through infant vocal exploration [6, 16]. To the best of our knowledge, the present paper presents the first large-scale empirical study systematically investigating the non-random occurrence of the three most salient phonatory protophone types of infancy, a study intended to provide perspective on the formation of vocal categories across the first year of life. Our analysis focused on vocants, squeals, and growls. Since vocants are the seeming default category (representing 80% or more of protophones), and since they manifest normal phonation, the type of phonation occurring in the vast majority of utterances in languages all over the world, our analysis focused on the clustering of protophones with non-normal phonation (squeals and growls) with respect to vocants. As seen in Fig 1c, more than 60% of infants exhibited significant clustering patterns on average across the ages for either squeals or growls. Moreover, every one of the 130 infants showed significant clustering of either squeals or growls in at least one recording. Across the six age groups, we observed significant clustering at all of them. Our findings offer robust empirical evidence supporting the idea that infants engage in systematic production of three vocal categories from the first months of life.

It was not unexpected that infants would show strong signs of clustering of squeals and growls in the age range from 3-4 months, because the idea of vocal play in infancy has been associated with stage models of vocal development that emphasize seeming vocal practice and vocal category repertoire “expansion” typically occurring during those months [6, 16]. But our empirical investigation did not provide support for the idea that 3-4 months of age is the predominant period of clustering. On the contrary, that age range showed the *lowest* proportion of significant clustering for both squeals and growls. The data in fact suggest that clustering was common at every age, with a tendency for the highest rates to occur from 5 months forward.

We are not, however, inclined to view this pattern of *age results* as offering the final word about vocal category clustering, if for no other reason, because there are many possible ways of coding phonatory types in infancy, and different approaches to coding could produce different age results. We adopted an efficient, simplifying approach, implementing real-time coding at the “utterance” level, restricting coders to three primary phonatory possibilities for each protophone utterance (vocant, squeal and growl). This approach made it possible to collect a large sample of data (assessing 1154 all-day recordings) with the resources available, but it clearly glossed over massive complexities. For example, *within* protophone utterances, more than one phonatory type often occurred, as illustrated in S4 Fig. a and b, and as a result, there is no absolute standard of correctness for categorization of such utterances either on auditory or acoustic grounds. Furthermore, even the 3 phonatory types could have been subcategorized, if there had been time and money to implement more extensive coding. The squeal type is manifest in at least two high-pitch vocal types (loft/falsetto and harsh) and the growl type in at least two non-high-pitch vocal types (pulse/fry and harsh). Vocants also are manifest in subcategories that include at least modal, tense and breathy types. Emphasis should be applied to the “at least” phrase in these depictions, because there are yet additional vocal regimes at stake—for example, “harsh” growls can include at least three vocal regimes (subharmonic, biphonation, and chaotic, see [54]). So while our coding system is systematic and yields reliable data, it may have oversimplified the vocal categories of human infants. The age pattern in Fig 1c that suggests more clustering beyond 5 months than before 5 months is thus subject to question because of the limits of our coding system.

These provisos in mind, we offer a few additional thoughts about the apparent age patterns reported above and their relations with speculations derived from stage models (especially [16]) that have influenced our expectations about when vocal play should be expected to occur. The fact that significant clustering was seen even in the youngest group in the present study was something of a surprise (although other research has also shown surprisingly early flexibility of protophone production [55]). We anticipated finding at least a significant increase in clustering from the first to the second age interval. The occurrence of significant clustering beginning at the very first age interval suggests that some mechanism of vocal exploration is present from the beginning of life, even before cortical control is thought to fully emerge. Prior to the descent of the larynx and expansion of the oral cavity, practice of vocal types may be focused on phonatory control categories. Whether additional protophone categories, including primitive supraglottal articulations (e.g., gooing) as well as raspberries, whispers, and yells, which become prominent at later stages are also show clustering practice is a matter for future research. Our simplified real-time coding scheme limits our ability to clarify all the possible patterns of vocal practice at present.

Another issue of concern is the difficulty of consistent coding in accord with our method.

Identifying vocal types in protophones produced by newborns clearly poses challenges beyond those requiring differentiation *among* the three phonatory types. Additional categories of infant sound making complicate the coding. For example, differentiating effort grunts (which often occur as a byproduct of movement, and which we exclude from the protophones by definition) and growls can be difficult due to their often shared characteristic of phonatory harshness. As infants get older, it appears most growls come to have longer durations, while effort grunts remain short, making it easier to recognize growls as distinct from grunts. During the newborn period, distinguishing between grunts and growls might then be viewed as resulting in an overestimation of growl quantities and thus tending to yield considerable apparent growl clustering at the earliest ages. The use of the term “grunts” in the vocal development literature sometimes to refer not to effort grunts but to quasivowels used to communicate assent toward the end of the first year [56] or simply to continue vocal conversation also suggests a complication with the counting of vocants (quasivowels being among them in our interpretation). These coding issues, along with others, clearly complicate workable interpretations of the early age patterns.

Another unexpected finding was the relatively stable significant proportions of growl clustering across the 6 age groups (Fig 1a), compared to higher proportions of squeal clustering at ages beyond 5 months (Fig 1a). The apparent differing developmental trajectories for growls and squeals might be attributed to some special characteristic of maturation of vocal fold control that might affect squeal and growl production differently across ages. Squeals, by definition, are produced in high pitch, whereas growls are produced with low pitch, and raising vs. lowering of pitch requires complex coordination of very different combinations of intrinsic and extrinsic muscles of the larynx [54] with different aerodynamic requirements, the development of which is not yet well understood. While the present coding results suggest infants do produce squeals from birth, as they grow and gain greater control of laryngeal muscles, perhaps they become able to exhibit more consistent production of squeal register, especially for the falsetto/loft regime.

On the other hand, growls typically occur at lower pitch and, it seems, often show harsh phonation that may require less fine glottal control than in falsetto/loft squealing. Perhaps, then, it is not particularly challenging for infants to produce harsh growls even from the earliest months, leading to somewhat similar proportions of clustering of growls across different ages. Squealing might in contrast grow in clustering across ages because the glottal control required for falsetto/loft may require maturation. These are of course speculations. Growling can also occur in non-harsh forms, with a pulse/fry regime. Does the pulse/fry regime require maturation similar to that for falsetto/loft? We do not know, and this supplies yet further reasons that our interpretation of the apparent age effects must remain uncertain.

Yet another issue worth considering is that inter-coder agreement was considerably higher for squeals than for growls. The segment-by-segment correlations in the agreement data for numbers of squeals within recordings showed 88% of pairwise comparisons were statistically significant, while 49% of pairwise comparisons were statistically significant for growls. Thus it seems possible that coders may not have been able to accurately capture developmental changes in growl production to as great an extent as for squeal production.

Even in the context of these difficulties of interpretation, our simplified approach to research on clustering of infant vocal types appears to reliably reflect the existence of three perceptually common protophone types that are often reported spontaneously by parents as forming features of infant repertoires. Squeals and growls represent salient departures from the default form of vocants that parents typically call vowel-like sounds. In the common parlance of many natural languages, squeals and growls are often represented with special terms [6]. Indeed, researchers of infant vocal development in the English-speaking world have adopted the terms squeal and growl directly *from* the common parlance, precisely because parents use them to describe salient infant vocal categories. Our inclination consequently is to take the data from the present real-time coding study seriously, at least to the extent that the data confirm the common parental report of the existence of these vocal categories in human infancy and help substantiate the impression that infants indeed practice these categories. We are also inclined to emphasize that the perceptions of human parents (as well as adults who do not have children, but often serve as caregivers and as member of our coding teams) are the natural gold standard of judgment about the nature and importance of infant vocalizations.

Thinking biologically, it is sensible to assume that human adults serve as the primary selection force on infant vocalization both in type and in quantity. If we are at some point to succeed in developing an automated system for categorization of human infant vocalizations, the standard of judgment for the success of that automated system will be the extent to which it can simulate the judgments of adult human listeners. So when we ponder on why there is so much infant vocal activity and so much clustering of vocal types within that activity, the biological perspective harkens back to the selective role of caregivers. The fitness signaling theory [57–59] suggests that ancient hominin caregivers noticed (and modern human caregivers notice) the occurrence of infant protophones and judge them in terms of the extent to which they indicate the wellness of infant vocalizers, providing a basis for selective investment in infants whose protophones are most indicative of fitness. Infants are not required to direct their vocalizations to caregivers (although it is important that they sometimes do so), because caregivers can notice protophones as indicating an infant is well and progressing normally toward vocal communication even if the infant is simply playing with sounds.

Perhaps one of the primary ways infants can reveal their wellness to caregivers is by producing protophone types in clusters that suggest practice with their spontaneously developed vocal categories. One might ask how competent parents could possibly fail to notice vocal clustering. Surely recognition of clustering of vocal types suggests infants are acquiring a system, admittedly a prelinguistic system, but a system of vocal categories under infant autonomous control. The seeming practice may even serve the purpose of confirming to infants themselves that they are on a path of increasing voluntary vocal control. Whether the infants realize it or not, that path reveals their possession of a capacity and an inclination not seen in any other ape, a capacity and inclination for voluntary manipulation of vocal categories, a capacity and inclination without which, it has been argued [59], language would be impossible.

## Acknowledgments

We would like to express our gratitude to the families whose infants participated in this research and to the graduate student coders in Memphis.

## Supporting information

Supporting information includes details on coding categories and coding agreement. It also includes waveforms (S1-S5_Waveforms) and spectrograms of each vocal category (S1-S5_Figs.) along with relevant details.

## Supporting Information

### Coding categories, definitions and coding criteria

The coding ignored vegetative sounds such as coughing, sneezing, and burping, none of which are treated as signals evolved as communications. It also ignored effort grunts, which we treat as artifacts of straining or movement. Coding of potentially communicative signals was conducted at the “utterance” level for all categories [1]. An utterance was defined as a breath group, where an inhalation was always taken to constitute an utterance boundary. Utterances could include silences (glottal holds or consonant-like closures, for example) during which no inhalation occurred. Also, a period without an inhalation could be judged, at the coder’s discretion, as an utterance boundary, if it included time enough for an inhalation, even though none was heard.

The real-time coding of utterances included 9 options for categorization of each utterance: vocant, squeal, growl, ingress, whisper, cry, whimper, laugh, and other (which included voiceless raspberries, voiced ones being categorized by their phonatory properties as vocant, squeal or growl). Ingress, whisper, laugh, and other constituted only ∼1% of the coded utterances, and were not included in the analyses for the present paper. Of note, surprisingly low rates of laughter have also been reported in prior work from our group, even in cases of laboratory recordings where parents play with their infants [2, 3]. Cries, whimpers, and laughs were also comparatively infrequent (constituting <14%) of utterances, with the three primary protophones (vocants, squeals, and growls) accounting for approximately ∼85% of all utterances across the first year. The criteria for categorization of cry and whimper utterances is explained in detail in a prior paper, along with acoustic specifications [4].

The present study focused on clustering of squeals and growls with regard to the very frequently occurring vocants, which constituted 74% of all utterances in the sample according to the coders, and more than 80% of the 3 phonatory protophone types. Growls in the present work, with its real-time coding, turned out to be about 7% of all utterances and squeals about 5%. This rate of squealing and growling is lower than has been found in studies using repeat-observation coding of laboratory recordings, e.g., 14% squeals and 17% growls in [3]. Our preferred interpretation of this discrepancy is that real-time coding (with only one opportunity for judgment per utterance) results in a tendency of coders to choose the “default” vocant category even more frequently than they do when listening to each utterance more than once. Thus, we presume that real-time coding tends to restrict the number of judgments of the non-default phonatory categories (squeal and growl) to particularly salient instances of their occurrence. We also think this restriction may result in a kind of recognition of clustering that may resemble the recognition by parents, who like the real-time coders, have only one opportunity with each infant utterance to gauge whether it represents a salient category.

Instructions to coders discouraged the inclusion in the coding of sounds that were so quiet or short in duration that they would be likely not noticed by caregivers. The basic approach to coding, includes the assumption that coders (almost all of whom are female in our studies) can and should listen and judge baby sounds the way caregivers do. After all, parents provide important influences on the infant in language adaptation, and if they could not make good judgments about the significance of infant vocalizations, they would surely be at a disadvantage in guiding their infants’ development. As a consequence, they should be at a disadvantage in promoting their own genes into subsequent generations, since their infants would presumably be at a disadvantage in survival and reproduction.

In the following sections, we provide definitions for vocants, squeals and growls. Buder et al. [5] and Oller [6] provide additional information regarding definitions of infant vocalizations (and particularly regarding the phonatory protophones) used in our laboratories. It is important to offer some explanation about the coding of the utterances because they are complex, yielding much better than chance agreement among coders, but also plenty of disagreement. There is considerable coding ambiguity, given that the categories are not fully discrete, but instead represent fuzzy classes with gradations among them, and often include within-utterance changes from one regime of phonation to another, yielding the basis for many coder disagreements. We reason that these disagreements are to be expected because the categories emerge, not from fixed innate inclinations, but from infant exploration within a complex space of vocal capabilities and tendencies.

All the figures presented in the Supporting Information were extracted from LENA recordings, and the spectrograms in the bottom panels of each figure are displayed at an 8 kHz range with a 30 Hz analysis bandwidth.

*Vocants:* The term “vocants” [7] in the present study includes both “quasivowels” and “full vowels” (also called “fully-resonant nuclei”, [6] ). Vocants are presumed precursors to mature vowels in natural languages. Quasivowels are produced with a vocal tract at rest, while full vowels are produced with a postured vocal tract [6]. Both quasivowels and full vowels are produced with normal phonation and fall within the infant’s habitual pitch range [5] which is typically 350-500 Hz [8]. The vocal “regime” of vocants is also often referred to as “modal” [5].

S1 Fig a and b provide oscillographic (time domain, top half of each panel) and spectrographic (spectral domain, bottom half of each panel) portrayals of two utterances judged as vocants from LENA recordings of 3-month-old infants. All the spectrographic displays in the figures in Supporting Information were produced from the original LENA recordings and display the full 8 kHz range available with a spectrographic bandwidth of 30 Hz, which allows for visualization of harmonics of the infant voices. As illustrated, the most prototypical vocants have clear harmonic structure with relatively little interharmonic noise. The utterances in the figure are both over a second in duration, and both are interpreted by coders as possessing at least two syllable prominences, but many vocants are much shorter, often 100 ms or less, and are often perceived as having only one syllable.

**S1 Fig.**
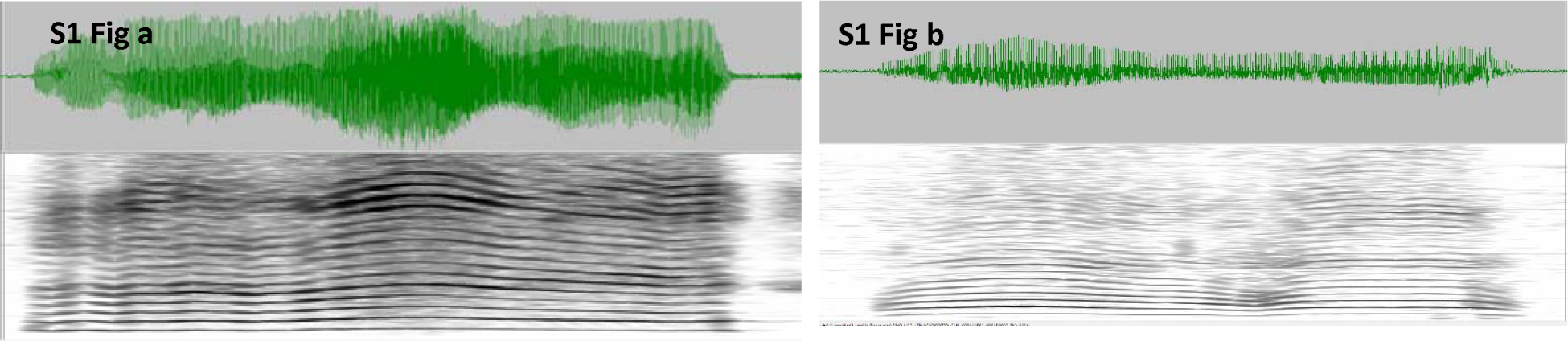
Display of vocant examples: a. A vocant from a 3-month-old infant, with a duration of display of 1269 ms. Notice the harmonics are clearly visible across the entire range of the utterance, with relatively little interharmonic noise. **b.** Another vocant from a different 3-month-old, with a duration of display of 1249 ms, again showing clear harmonics and little interharmonic noise.

*Growls:* Growls tend to have lower pitch, sometimes much lower pitch than vocants. An utterance with pitch substantially higher than typically accompanies the modal regime in any infant is not allowed to be categorized as a growl within our scheme. The vocal regimes of growls often yield an impression of substantial harshness as corresponds to the majority of the utterance represented in S2 Fig a. An acoustic manifestation of the harshness is the considerable amount of interharmonic noise, which can be the product of chaotic, subharmonic or biphonation regimes [5]. S2 Fig b presents another way that growls can manifest, namely with extremely low pitch, where individual pitch periods can be discerned in the time domain display, and where the ear perceives a “zipper” quality attributable the listener’s being able to hear the individual pitch pulses. This regime is thus often called “pulse” although linguists tend to term it “vocal fry” or “creaky voice.”

**S2 Fig.**
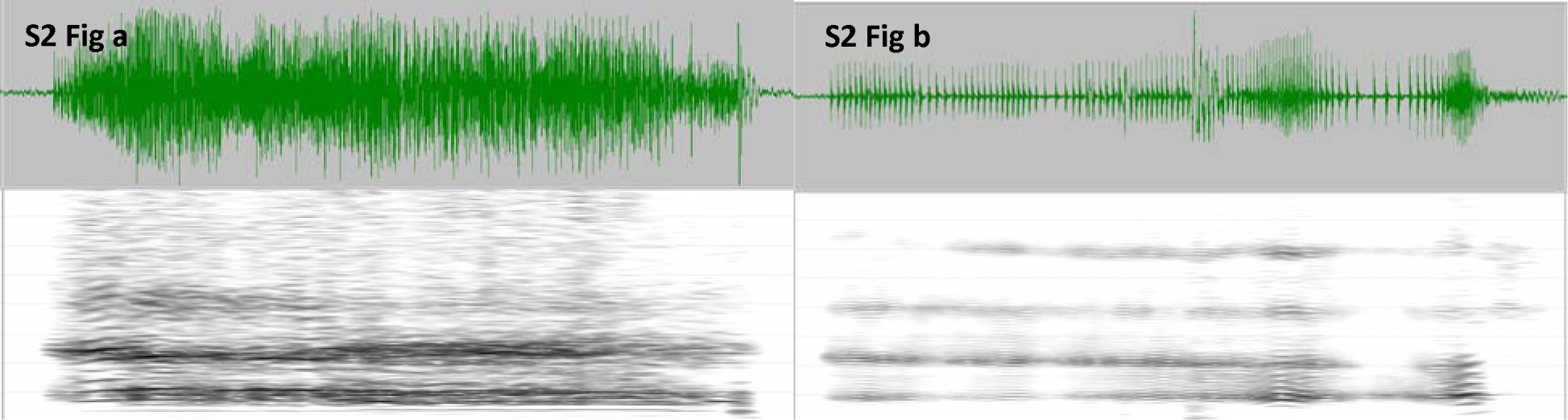
Display of growl examples: a. A harsh growl from a 3-month-old infant, with a duration of display of 1215 ms. During this utterance there is considerable interharmonic noise, which contributes to the perception of phonatory harshness. The first fifth of the utterance does not show the harshness of the rest of the utterance. **b.** A second growl from a different 3-month-old, with a duration of display of 1271 ms, is very different from the one in panel a, because this one has the phonatory property of pulse (or vocal fry) throughout. Notice that harmonics in 2b are very narrowly spaced and that individual pitch periods can easily be discerned in the time domain display at the top. In both 2a and 2b coders tend to hear the pitch as being considerably lower than in vocants.

*Squeals:* S3 Fig a and b show squeals from two 3-month-old infants. The vocal quality in S3 Fig a corresponds to falsetto (or “loft”) phonation, as indicated by very widely spaced harmonics compared with the vocants of 1a and 1b. S3 Fig b is referred to in our laboratory as a harsh squeal, which has periods of relatively pure falsetto with little noise between harmonics, but the most salient parts of the utterance are very harsh and high in pitch. There are even periods of normal phonation during this long and complex utterance. The training in our laboratories encourages coders in the forced choice real-time coding to categorize utterances with such complexities in accord with their most salient features. In the case of 3b, coders tend strongly to code the utterance as a squeal.

**S3 Fig.**
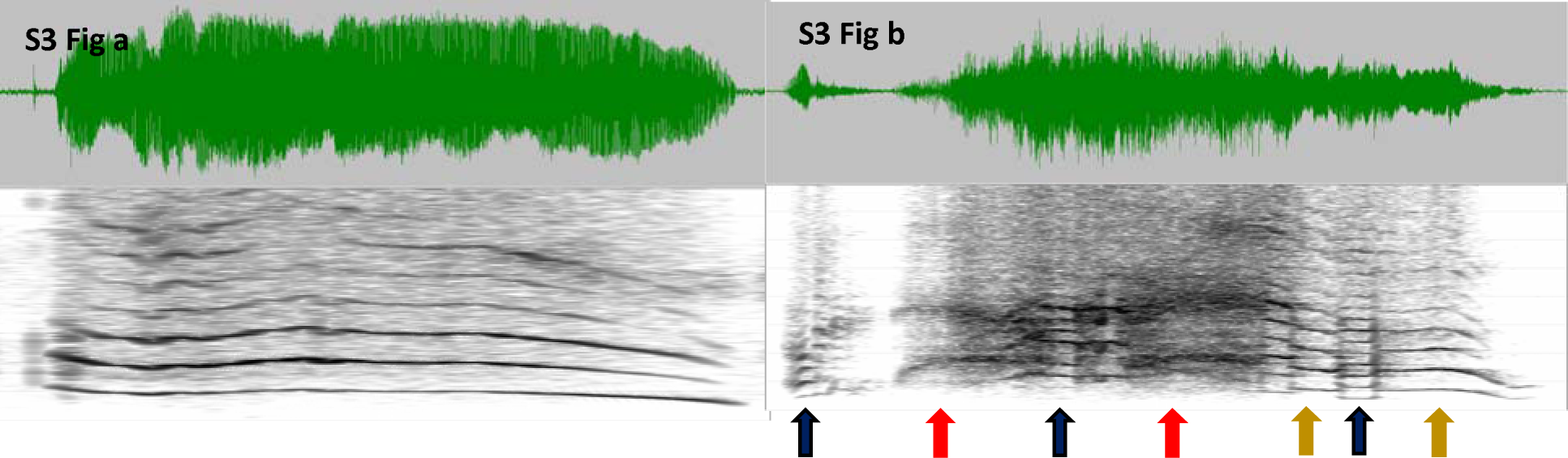
Display of squeal examples: a. A squeal from a 3-month-old infant, with a duration of display of 1270 ms. Notice the harmonics that are widely spaced, corresponding to very high pitch accompanying the falsetto or “loft” regime and like the prior vocant examples, showing relatively little interharmonic noise. **b.** A much more complicated squeal from a different 3- month-old, with a duration of display of 3141 ms, showing clear widely spaced harmonics with relatively little noise between harmonics only during two periods designated by gold arrows. The red arrows indicate periods of noisy, harsh squeal phonation, with a great deal of noise, perceived when isolated as very high in pitch. Two brief periods of relatively modal phonation, with more narrowly spaced harmonics are designated by black arrows, and if isolated, sound like vocants. There is a brief period shortly after the initial period of normal phonation where there is an apparent glottal hold, resulting in very little sound. The utterance as a whole was unambiguously categorized by coders as a squeal, because the most salient auditory features of the utterance were deemed to include very high pitch. We encouraged coders to consider any salient period of squeal or growl phonation during an utterance as reason to avoid the vocant category, although there are plenty of cases where low intensity or short squeal or growl phonatory patterns are found in an utterance that coders judge as vocant.

*Utterances with notable regime shifts yielding ambiguity of judgments:* The coding task required a forced choice at the utterance level, and consequently coders were not uncommonly faced with utterances that had multiple possible interpretations, and sometimes, unlike the utterance in S3 Fig b (where the tendency was strong to choose squeal even given the regime complexity), there was more basis for coder disagreement. S4 Fig a and b provide good examples of hard-to-judge utterances. In both 4a and 4b we see multiple regime shifts. Coders are variable in their decisions about both these utterances, sometimes choosing squeal and sometimes growl, but never vocant.

**S4 Fig.**
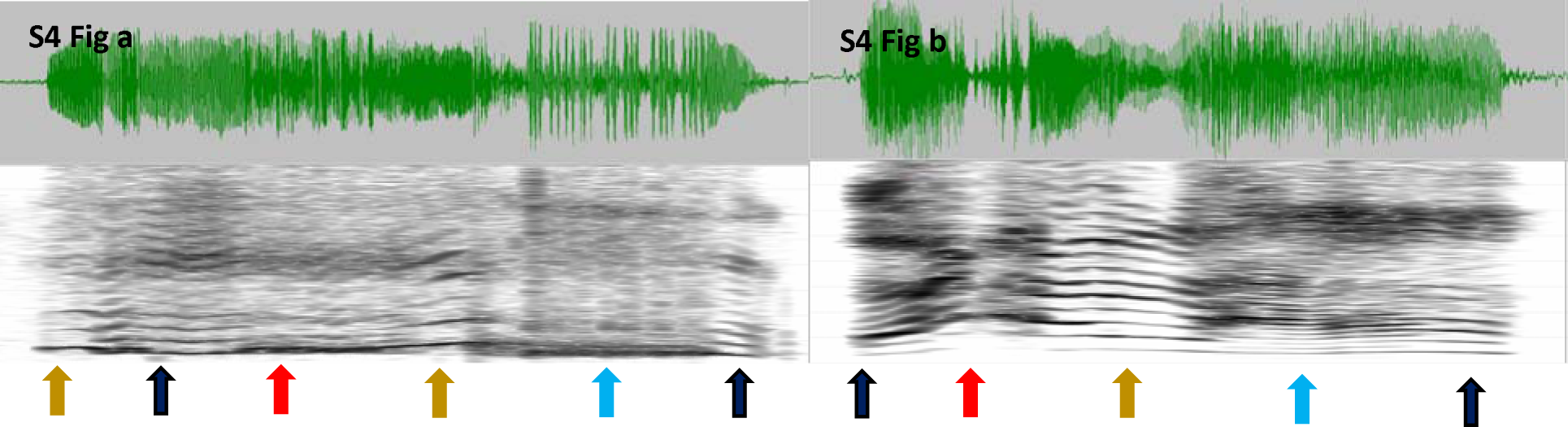
Display of complex utterances with uncertain interpretation: a. A complex utterance from a 6-month-old infant, with a duration of display of 1384 ms. As before, gold arrows point to periods of loft phonation, red to periods of high-pitched harsh phonation, and black to periods of normal phonation. Blue arrows correspond to harsh growl phonation. This is an utterance that is variably coded as squeal or growl. **b.** Again, a complex utterance, 863 ms duration of display, from a different 6-month-old infant, shows several regime shifts and is judged variably as squeal or growl. These utterances would never be judged as vocants in the OLL.

*Spectrographic illustrations of clustering:* In S5 Fig a and b we present examples of the clustering phenomenon. In S5 Fig a, 7 squeals occur in a row, in S5 Fig b, 6 vocants in a row. In these cases, the utterances occur in a repetitive pattern for squeals in one case and vocants in the other. In the present study, clustering did not require repetition, because we counted the number of utterances of each phonatory type within 5-minute segments of recording, and then compared 5-minute segments for the numbers of each phonatory type across the segments drawn from each all-day recording. Repetition could result in significant clustering in our analysis, but clustering could also occur when the rate of squeals or growls was higher in some segments than in others in spite of the occurrence of mixing with other vocal types within the segment.

**S5 Fig.**
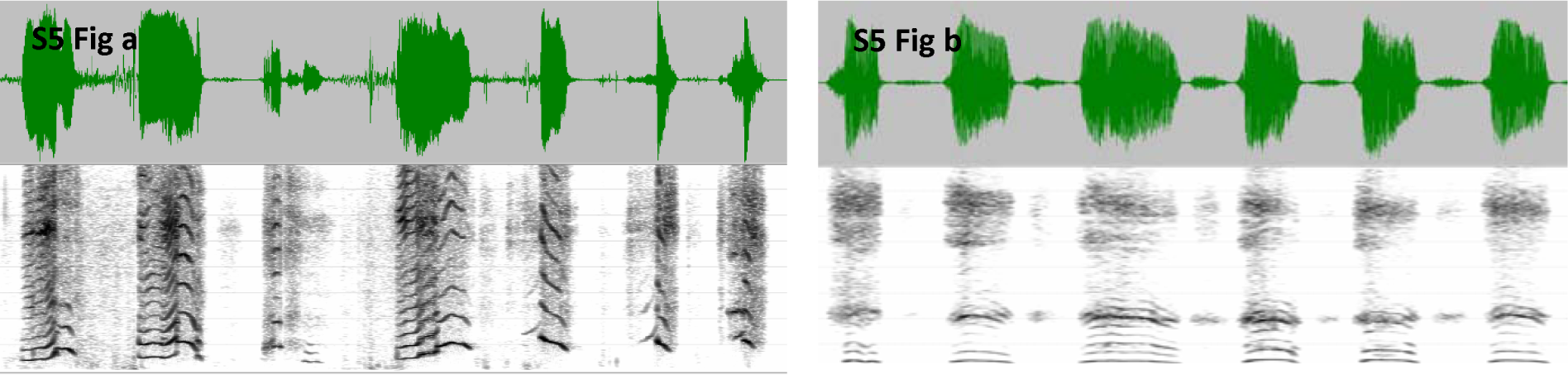
Clustering Acoustic Display: a. A string of 7 utterances, all judged as squeals, from a 6-month-old infant. Notice the brief pauses between utterances, and notice that each utterance has a notable period of high pitch (falsetto), which forces the squeal judgment in accord with our coding criteria, in spite of the fact that modal phonation also occurs in several of the utterances. The duration of display is 6308 ms. **b.** From the same infant in panel a, we see a sequence of 6 vocants from the same recording day, all with clear modal phonation, and in this case there are visible inhalations between the utterances in most cases.

### Illustrations of the application of Fisher’s exact test

S1 Tables a and b present examples from the real data of application of the Fisher’s exact test for two different recordings from two of the infants where the comparison is between squeal and vocant counts. Note that in S1a there are 14 surviving segments out of the 21 selected from each recording (segments having at least one protophone of each of the relevant types, segments not having more than 5 cries or whimpers, and segments where the infant was not asleep), while in S1b there are 12. In both cases the Fisher’s exact test yielded a significant result (*p* < .001), indicating that squeals and vocants did not distribute randomly with respect to each other. Notice for the example in S1 Table a that in segments 1, 4, 19 and 20, vocants dominated, while in 16 through 18 squeals dominated, suggesting a period of squeal practice/vocal play during segments 16 and 18. Similarly in S1 Table b, the infant shows clustering of squeals in segments 18 through 20. S1 Table c shows no statistically significant pattern of clustering of squeals.

**S1 Table:**
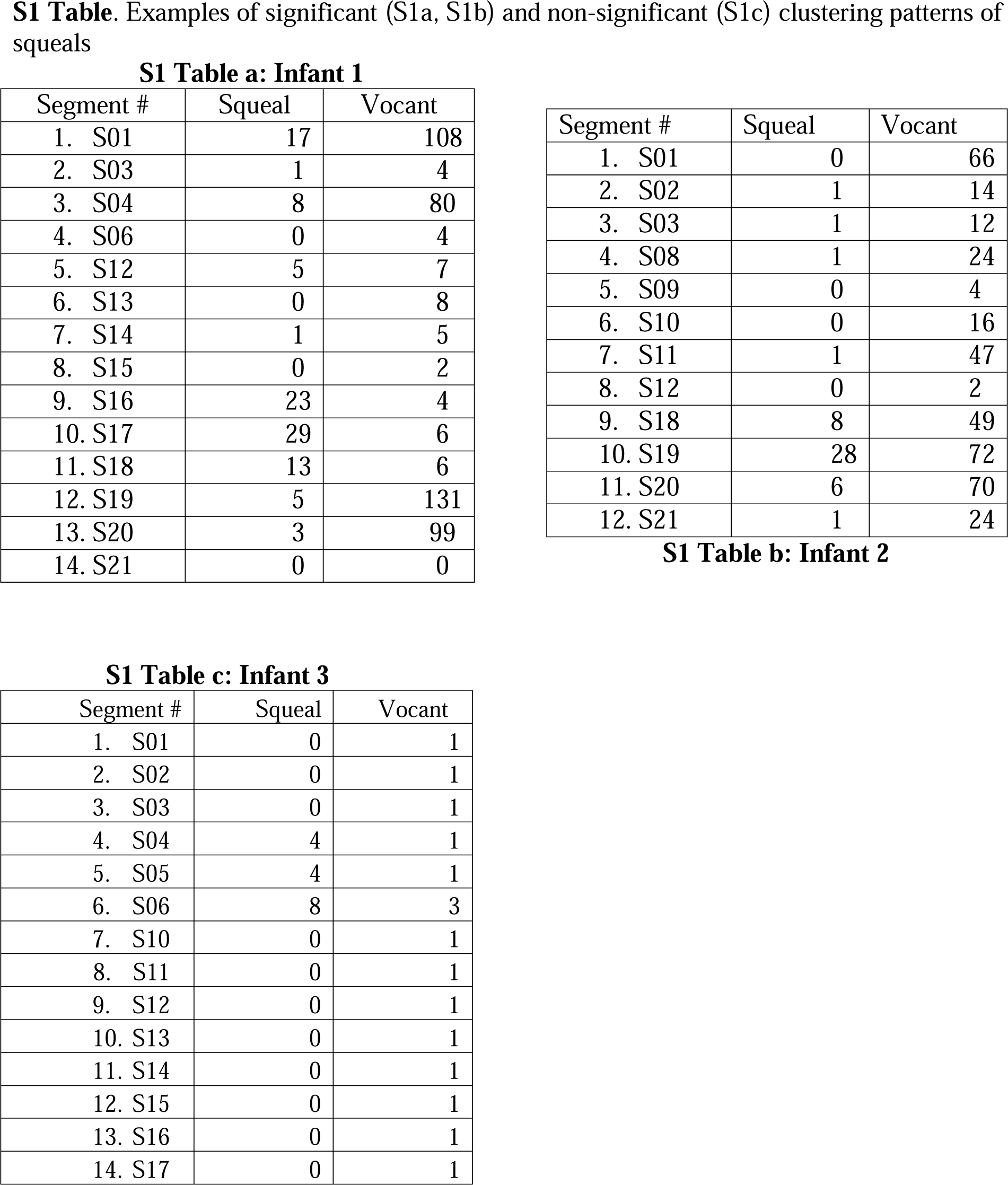
**a.** Numbers of squeals coded in 5-minute recording segments surviving the analysis filter (which excluded sleep segments and segments with 5 or more cries or whimpers) are contrasted with numbers of vocants in the same segments for one infant’s all-day recording. Note that some segments are missing in the tables, because they did not survive the filtering. The numbering reflects the sequential order of the surviving segments. It is clear that squealing tended to occur non-randomly with respect to vocants. **b.** In a recording of an additional infant, the sharp tendency for squeals to occur non-randomly with respect to vocants across surviving segments is also obvious. **c**. In the case of another infant’s recording, the Fisher’s exact test did not yield a significant result (*p* = .264).

S2 Tables a and b present an example of application of the Fisher’s exact test where the comparison is between growl and vocant counts. In S2 Table a, there were 19 surviving segments, where segments 5, 14 and 21 showed significant growl activity with respect to vocant activity, but not in any of the other segments. Similarly, in S2 Table b, where there were 18 surviving segments, there was high growl activity with respect to vocant activity in segments 8 and 21, for example.

**S2 Table:**
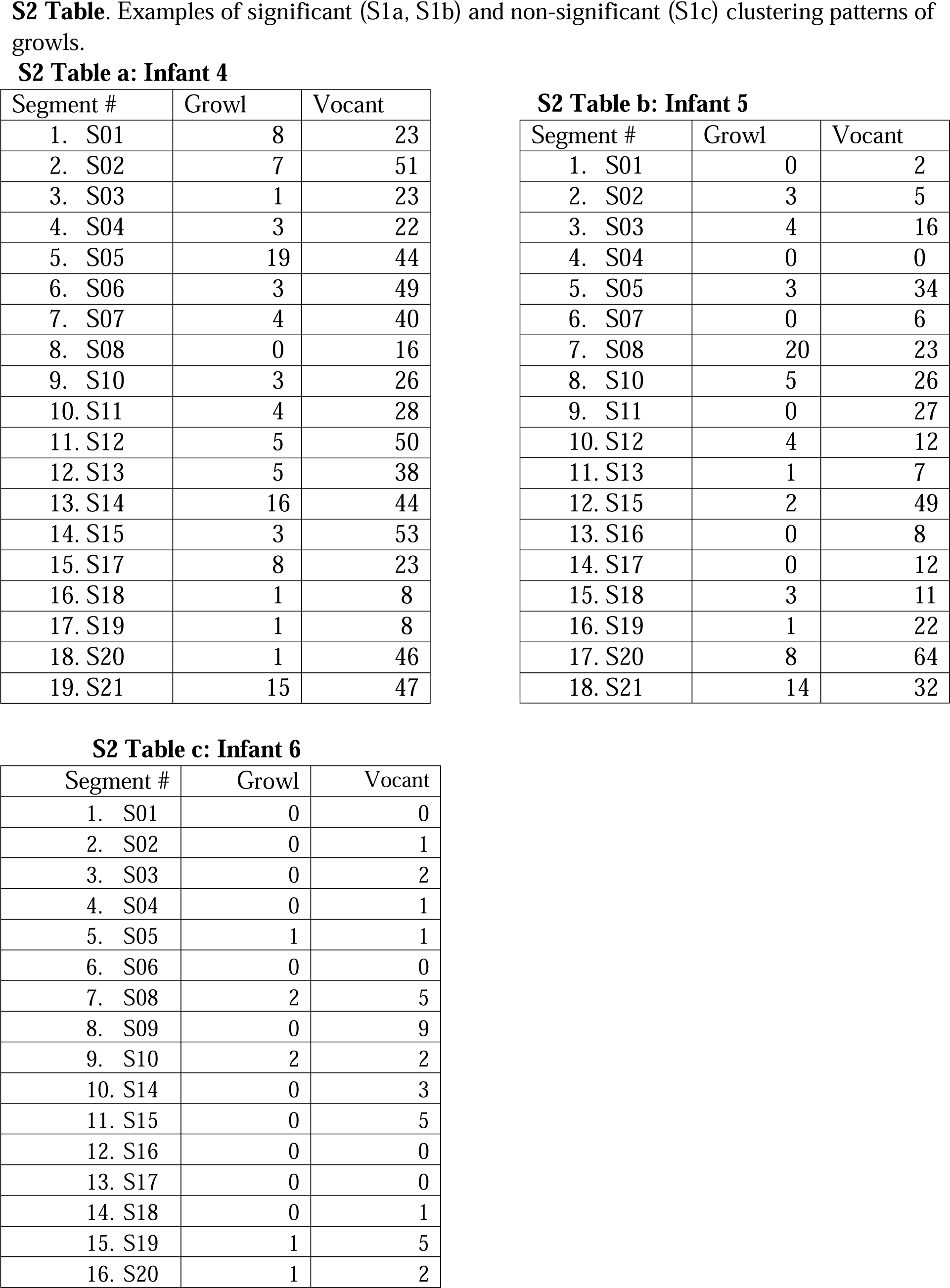
**a.** Numbers of growls coded in 5-minute recording segments surviving the analysis filter are contrasted with numbers of vocants in the same segments for an all-day recording from infant 4. It is clear that growling tended to occur non-randomly with respect to vocants. **b.** In a recording of infant 5, the sharp tendency for growls to occur non-randomly with respect to vocants across surviving segments is also obvious. **c.** In a recording of infant 6, on the other hand, the result was not statistically significant, although there were some growling sounds produced across the segments (*p* = .337).

Again, for both infant 4 and infant 5, the Fisher’s exact test yielded a significant result (*p* < .001), indicating that growls and vocants did not distribute randomly with respect to each other. S2 Table c shows no statistically significant pattern of growl clustering.

### Coder agreement

#### Results from correlational analyses illustrating statistically significant coder agreement

Providing a basis for determining the reliability of the coding of the three protophone types, 21 five-minute segments randomly selected from 9 day-long recordings (a training and agreement recording set) were coded independently by many individuals during the training period. The data from this training dataset supply a basis for an extensive evaluation of coder agreement, where all the coders who contributed to the Results of the present study participated, and where the agreement recordings were from the same Marcus Autism Center database as the recordings used in the main study.

All 21 segments of some of the 9 recordings were coded independently by all of the 36 coders of the teams that coded the data reported in Results. There were additional coders from related OLL projects who had been trained the same way who also coded some or all of the 9 recordings. Thus, there was a variable number of individuals who coded each of the 9 recordings, with an average of 32.1 coders per recording (range 20-48). These agreement recordings were essentially identical to those of the data from the 312 infants described in Methods of the main text. The recordings were also obtained with the LENA system during the same time period of the recordings from the 312 infants of the broader set evaluated by the OLL, from infants recruited at the Marcus Autism Center in the same way, and meeting the same inclusionary and exclusionary requirements.

An additional agreement analysis is made possible by a dataset involving 9 of the coding team members who contributed to the Results. Each of these coding team members was assigned, after each one of them completed coding all the recordings on at least 4 infants to independently code semi-randomly selected 5-minute segments that had previously been coded by one of the other team members. This second dataset provided another opportunity to demonstrate that coders agreed substantially on the categorization of squeals, growls, and vocants.

But before considering the statistically significant agreement results, it is important to acknowledge that there was considerable variation among the coders in their counts of the three protophone types within recordings. This variation indicated, as was clear during the training and in prior research with such data [3], that coders often disagreed about categorizations of squeals, growls and vocants. A simple measure of the extent of this disagreement can be obtained from the coefficient of variation (CoV, ratio of standard deviation to mean) across coders for each of the 9 recordings where multiple individuals coded the same 21 segments independently. The mean CoV for vocants was 0.21 across the 9 recordings, suggesting a relatively narrow range of vocant counts across coders. Squeals and growls showed a much larger range, with mean CoVs of 0.59 and 0.96 respectively. Thus, the data for the present study are clearly subject to substantial differences among coders in how they made decisions, with the acoustically analyzed examples presented above hopefully supplying perspective on why we assert that such disagreements are to be expected. The higher CoV for growls is an indication that coder agreement was lower for growls than for squeals.

The analysis conducted on clustering and presented in Results can be justified by its much higher than chance level agreement among coders on the segment-by-segment counts of the protophone categories. We evaluated correlations among coders on squeals, segment-by-segment within recordings, as well as correlations among coders on growls. Spearman rank order correlations were computed on the 21 segments for each recording in this agreement set, first on the numbers of squeals and then on the numbers of growls classified by each coder in all the available pairwise comparisons of coders for each recording from the agreement set. In this way we determined the extent to which coding from each individual matched that of each other individual coder with regard to numbers of squeals or growls, 5-minute segment by 5-minute segment, across all 21 segments for each recording (there were 9802 individual correlations).

Then we conducted that same analysis on the 5-minute segments of the second agreement set.

The analysis on the first set yielded many statistically significant correlations (for each pairwise comparison, N = 21, and for ρ *>* .433, *p <* .05). An average of 88% of pairwise comparisons produced statistically significant correlations for the 9 recordings on squeals; for growls, 45% of pairwise comparisons were statistically significant. The mean Spearman correlation across the 9 recordings for squeals was ρ = .71 and for growls ρ = .42.

To demonstrate that this degree of agreement among coders is very much greater than chance, we bootstrapped randomizations of the observed counts for all the coders for all the segments within each recording, using two of the 9 recordings for the randomization tests. The randomizations produced much smaller percentages of significant correlations for the bootstrap coders than the real coders, averaging only 6% of significant pairwise comparisons for the bootstraps on both squeals and growls, with upper bounds of their 95% confidence intervals at 7%. The lowest percentage of significant correlations between real coders for the pairwise comparisons for any of the 9 recordings was 40% for squeals and 27% for growls, far higher than the upper bounds of the significant correlations for the bootstrapped randomization tests.

In addition, data from the second agreement study were analyzed. The agreement data were obtained near the end of the data collection from 40 of the infants who were coded by the Memphis team. Among the 36 trained coders, 10 individuals coded the 40 infants, and 9 of those coders were available to be assigned for agreement coding. We semi-randomly selected 523 five-minute segments from the 21 coded for each of the recordings and assigned each of these segments for blind recoding by an agreement coder, not the individual who had originally coded the segment. We balanced the assignments, to the extent possible, such that agreement coders were assigned segments pertaining to as many of the age categories and infants as possible given the coding time available for each individual—we succeeded in assigning each coder to segments from at least 5 of the 6 ages (mean = 5.67) and at least 19 of the 40 infants (mean = 20.67). The number of segments recoded by the agreement coders ranged from 19 to 119. Data on agreement for protophones, cries and laughs from this second agreement set were presented in a prior paper [2]. The average correlations for squeals and growls are presented here for the first time. The agreement levels between original and agreement coders were higher than in the case of the correlations among coders of the 9 training recordings. The mean across the nine coders on the second agreement set for squeals was ρ = .79 and for growls, ρ = .65). All but one of the individual 18 correlations was statistically significant. The higher correlations on the second agreement set appear to support our assumption that coding agreement improved with experience.

The fact that the inter-coder correlations significantly exceeded chance proves that the coders detected a reliable “signal” for squeals and growls in the data. That the correlations were not high by traditional standards does not invalidate the comparisons presented in Results. Low coder agreement, as long as it is statistically significant, has an effect on group comparisons (in this case, across-session comparisons to determine possible clustering) that limits the possibility of detecting the sought after effect because the low coder agreement acts as noise and can produce Type II error. But if significant effects (in this case, clustering effects) are detected in spite of the noise, there is no reason to doubt the outcomes—the effects can be said to have been robust enough to have significantly exceeded the noise.

### Results from permutation tests supporting intercoder agreement

Figures in this section present results from permutation tests [9] using all of the nine agreement recordings from the training period, aggregated to test for coder variation. We resampled subsets of the (at most) 48 coders’ clustering results (from one to 48) without replacement 5,000 times and calculated the proportion of all nine recordings that showed significant clustering patterns for squeals and growls (panels A and B in S6 Fig) with respect to vocants to determine the empirical distribution of the significant clustering that occurred according to the Fisher’s test under the null hypothesis that coder identity had no effect on the clustering outcomes. For C, the clustering patterns for either squeals or growls were taken into account, yielding the highest proportion of clustering for the three panels.

**S6 Fig.**
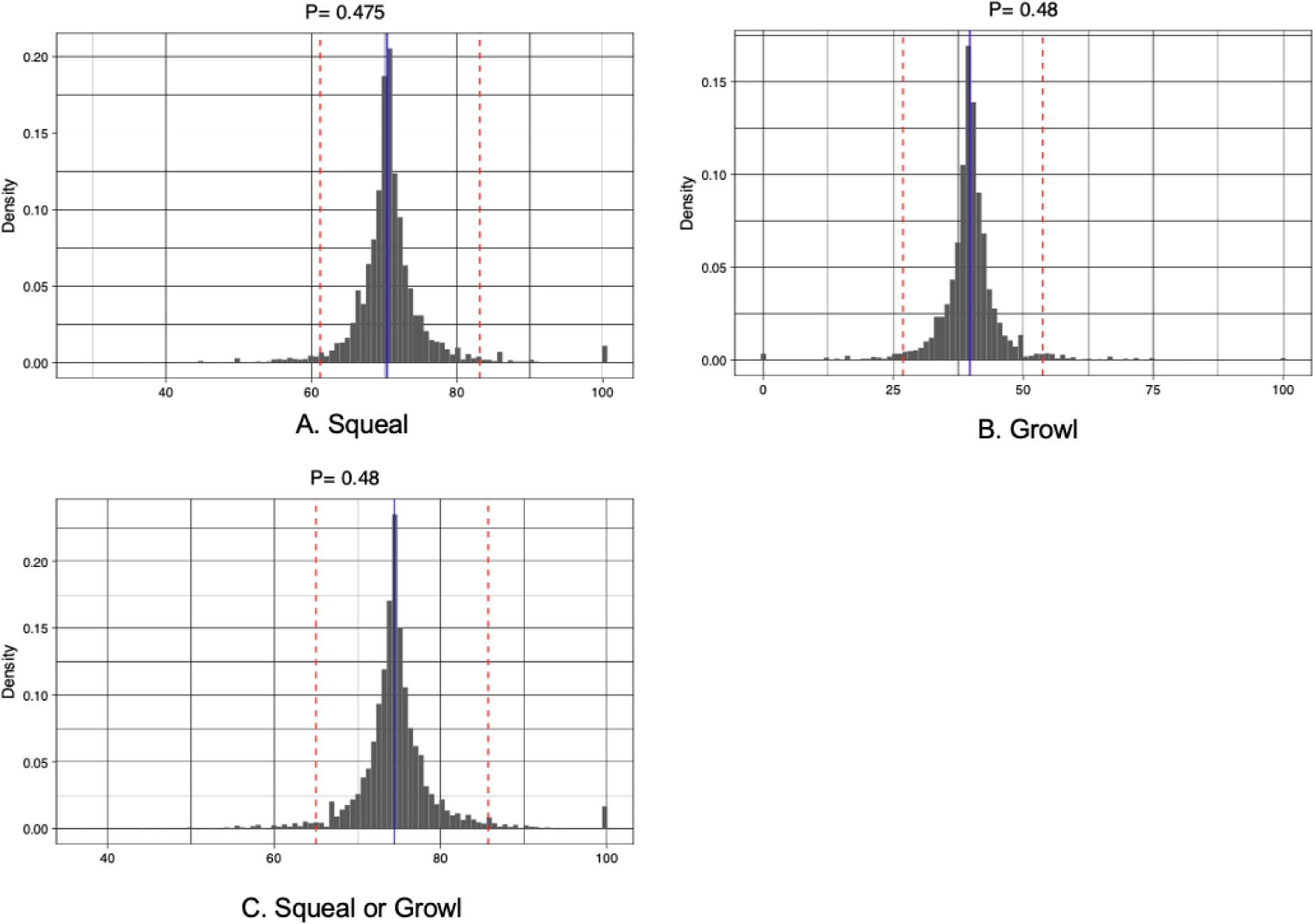
Permutation results for all agreement recordings.

For the three panels of S6 Fig, the X-axis represents the proportion of recordings that showed significant clustering on the nine recordings. The y-axis represents the probability density for each proportion of the nine recordings that were found to have significant clustering. The blue line represents the observed mean of the actual data (with no permutation) from all the coders across all nine recordings. The red dotted lines represent the 95% confidence intervals for the distribution of proportions of recordings that showed significant clustering resulting from the permutation test. The fact that all the blue lines fall deeply within the area (and approximately on the mode) between the red dotted lines confirms that differences between coders, although present (as indicated by coefficients of variation across coders across the 9 recordings), had little effect on the significant clustering findings for these recordings. If there had been little or no agreement among the coders, the blue lines would have fallen outside the 95% confidence interval as indicated by the dotted red lines. Thus, these findings supplied no evidence that coder differences significantly skewed the clustering results of the agreement recordings.

In separate analyses, the nine recordings were evaluated individually to produce the same kinds of permutation tests for squeals and for growls as well as for squeals or growls as shown in S6 Fig, although the number of individuals who had coded each of the recordings ranged from 20 to 48. Thus each permutation test for an individual recording was conducted just for the number of individuals who had coded all 21 segments from that recording. These analyses revealed very distinct distributions for different recordings, and even within recordings, often very different distributions for squeals and growls. For example, for recording 2 (S7 Fig) the distribution of coders’ proportions of significant clustering findings for growls (a very low proportion) scarcely overlapped with the distribution of coders’ proportions of significant clustering findings for squeals (a high proportion), and the confidence intervals were widely separated. In recording 9 (S8 Fig) the pattern showed very few significant findings of clustering for either squeals or growls. In recording 1 (S9 Fig) the distributions showed significant clustering for both squeals and growls, although the clustering pattern was found by the coders to be stronger for squeals. In recording 6 (S10 Fig) there was total coder agreement that squeals were clustered (and that is why the panels for squeals and squeals or growls are blanked out), while the distribution for coder findings on growls centered around 50% significant clustering. Across the nine recordings, the distributions strongly indicated that, as a group, the coders agreed on whether either squeals or growls or both showed clustering. As in the case of the aggregated analysis depicted in S6 Fig, if there had been little or no agreement among the coders, the blue lines for S7-S10 Figs could indeed have fallen outside the 95% confidence interval as indicated by the dotted red lines. But that did not happen in any case across the nine recordings, because the coder agreement was too high.

**S7 Fig.**
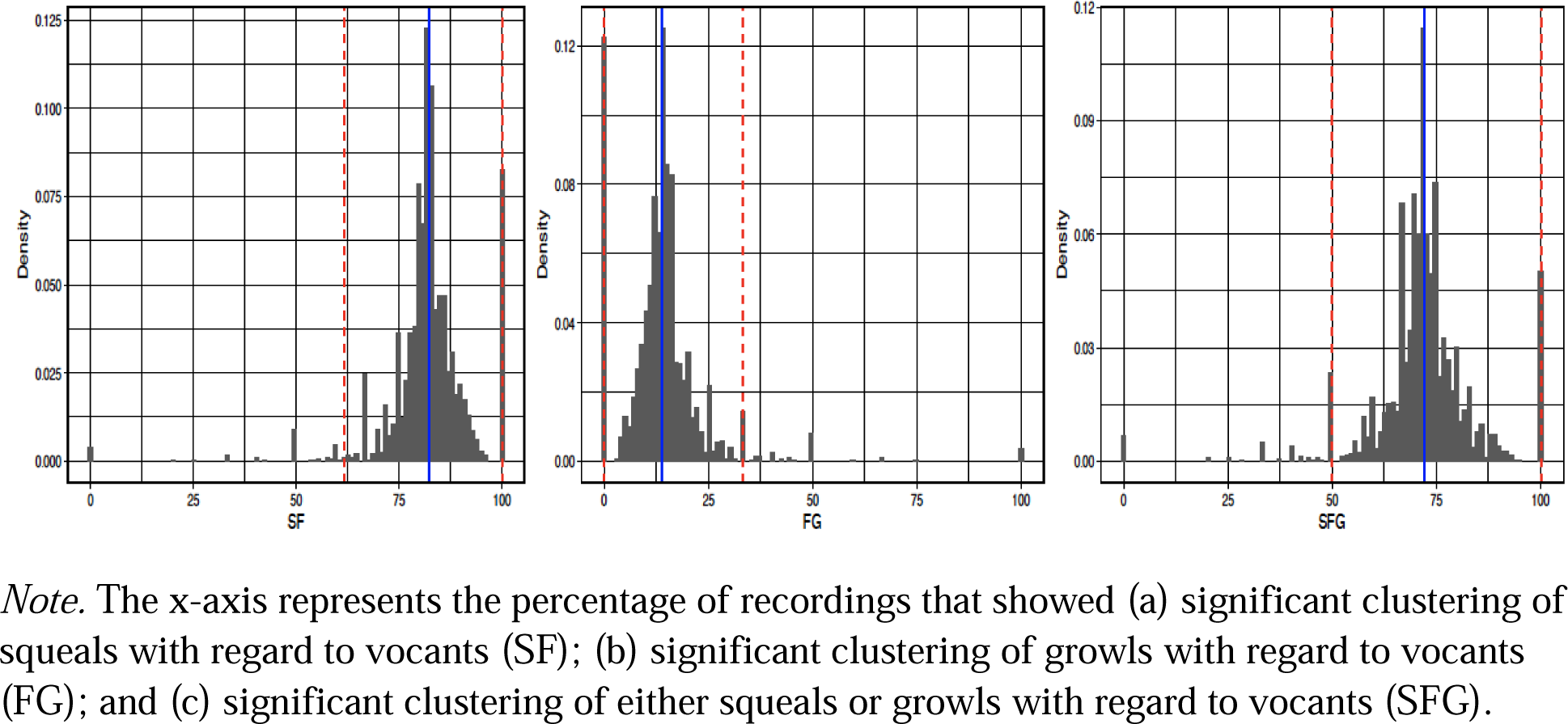
Permutation results for Recording 2. *Note.* The x-axis represents the percentage of recordings that showed (a) significant clustering of squeals with regard to vocants (SF); (b) significant clustering of growls with regard to vocants (FG); and (c) significant clustering of either squeals or growls with regard to vocants (SFG).

**S8 Fig.**
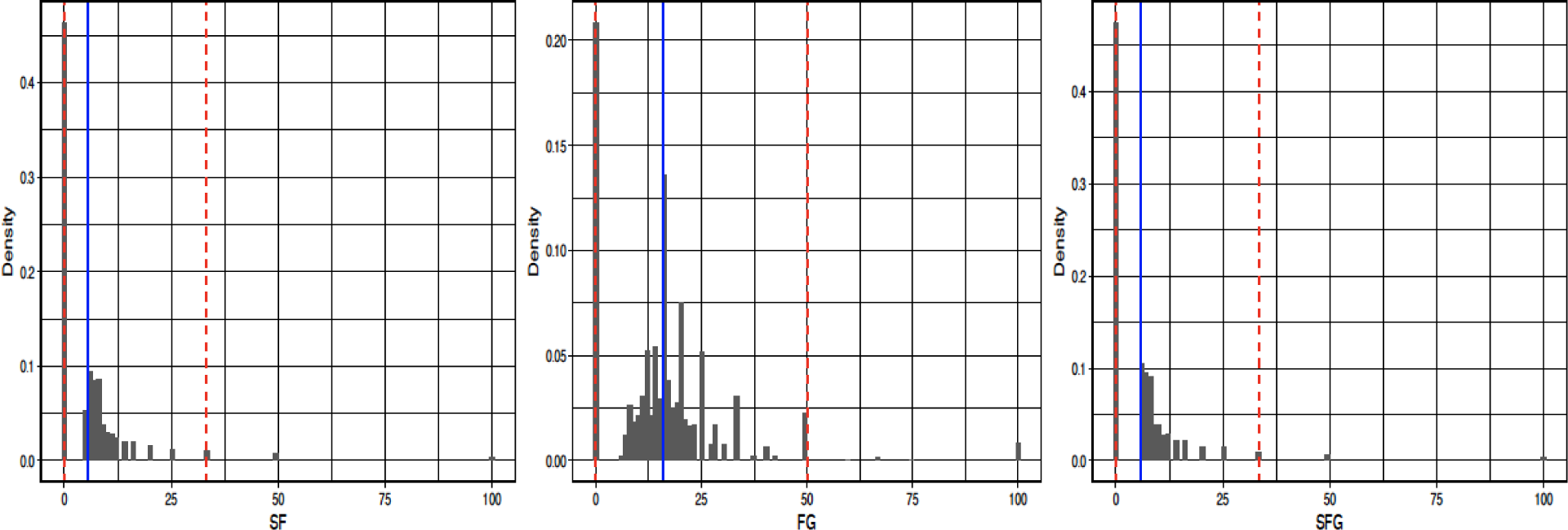
**Permutation results for Recording 9.**

**S9 Fig.**
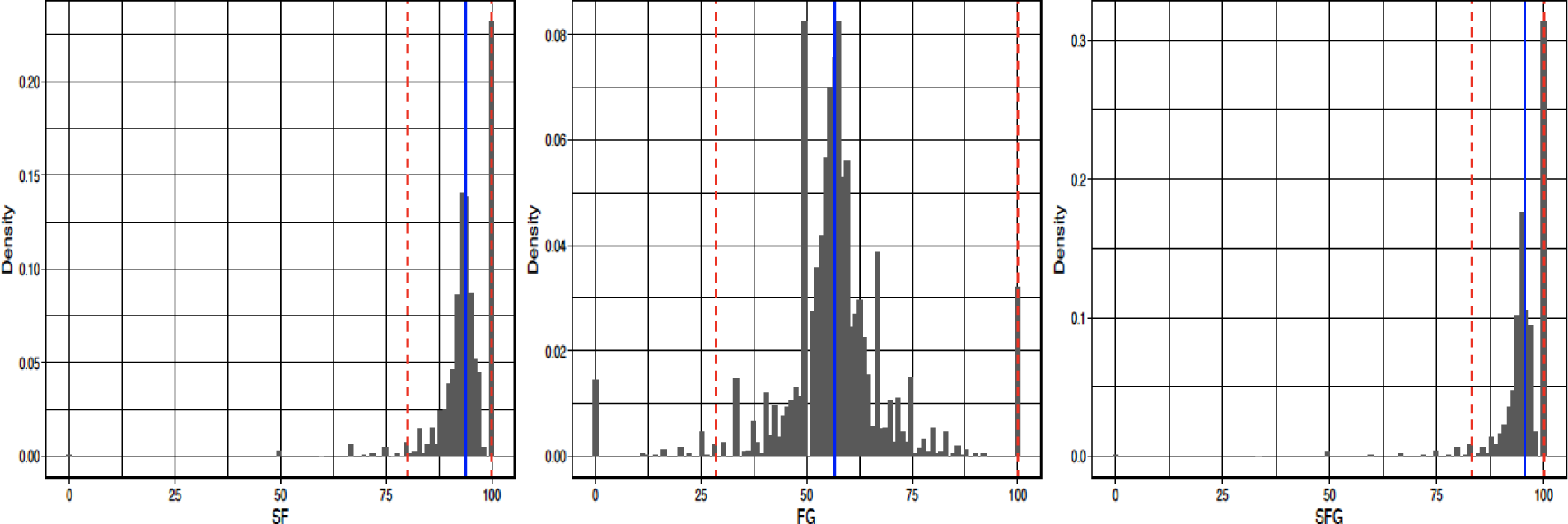
**Permutation results for Recording 1.**

**S10 Figure.**
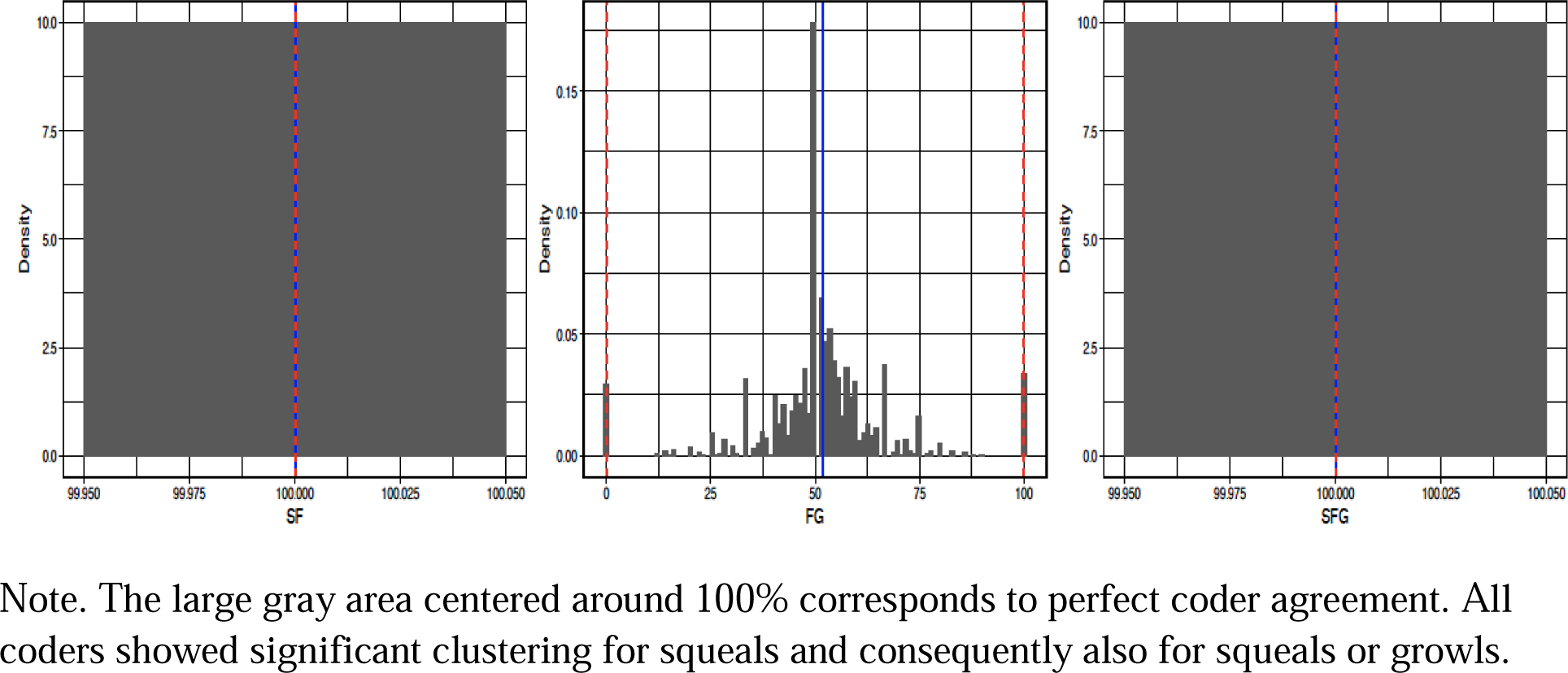
Permutation results for Recording 6. Note. The large gray area centered around 100% corresponds to perfect coder agreement. All coders showed significant clustering for squeals and consequently also for squeals or growls.

The conclusion of these agreement analyses is that, while there was considerable inter- coder variation, the overall agreement on coding was strong and gives reason for confidence in the clustering results reported in the main text.

